# PROSSTT: probabilistic simulation of single-cell RNA-seq data for complex differentiation processes

**DOI:** 10.1101/256941

**Authors:** Nikolaos Papadopoulos, R. Gonzalo Parra, Johannes Söding

**Affiliations:** Quantitative and Computational Biology, Max Planck Institute for Biophysical Chemistry, Göttingen, 37077, Germany

## Abstract

**Background:** Single-cell RNA sequencing (scRNA-seq) is an enabling technology for the study of cellular differentiation and heterogeneity. From snapshots of the transcriptomic profiles of differentiating single cells, the cellular lineage tree that leads from a progenitor population to multiple types of differentiated cells can be derived. The underlying lineage trees of most published datasets are linear or have a single branchpoint, but many studies with more complex lineage trees will soon become available. To test and further develop tools for lineage tree reconstruction, we need test datasets with known trees.

**Results:** PROSSTT can simulate scRNA-seq datasets for differentiation processes with lineage trees of any desired complexity, noise level, noise model, and size. PROSSTT also provides scripts to quantify the quality of predicted lineage trees.

**Availability:** https://github.com/soedinglab/prosstt

## 1 Introduction

Recent advances in single-cell RNA sequencing (scRNA-seq) (Macosko, 2015; Klein *et al.*, 2015) make it possible to generate expression profiles for thousands of cells. Applications to study cellular heterogeneity are challenging traditional definitions of cell types (Trapnell, 2015) and techniques are being scaled up for the Human Cell Atlas Project (Regev *et al.*, 2017). Single-cell transcriptomics is also pushing the study of cellular differentiation to a new, quantitative level. For example cellular lineage trees can be reconstructed from snapshots of single-cell transcriptomes of cells caught at various stages of their differentiation process. Organoids serve as particularly attractive, simple model systems (Camp *et al.*, 2017). The change in gene expression along the reconstructed trees gives us unprecedented, time-resolved data to quantitatively investigate the gene regulatory processes underlying cellular development.

Various tree inference methods have been developed (Rostom *et al.*, 2017). They order cells according to their pseudotime, i.e., to their progression along a developmental pathway. The progenitor population is at the root of the lineage tree, branchpoints in the tree represent cell fate decisions, and endpoints represent differentiated cell states.

Lineage tree inference algorithms are indispensable for the analysis of scRNA-seq data. As more and more complex processes are investigated, there will be a need to derive lineage trees of topologies more complex than linear or singly-branched ones. Also, with various methods already published and more being developed, the need to quantify method performance is becoming more pressing. With the available data, assessing method performance is challenging as there are no datasets with known ground truth, i.e. data with known intrinsic developmental time and cell identity.

To address these needs we developed PROSSTT (PRObabilistic Simulation of Single-cell RNA-seq Tree-like Topologies), a python package for simulating realistic scRNA-seq datasets of differentiating cells.

## 2 Model

PROSSTT generates simulated scRNA-seq datasets in four steps:

1. **Generate tree:** The topology of the lineage tree (number of branches, connectivity) and the length of each branch are read in or, alternatively, sampled. The integer branch lengths give the number of steps of the random walk (see next point) and correspond to the pseudotime duration (Fig. 1A (inset)).
2. **Simulate average gene expression along tree:** Gene expression levels are linear mixtures of a small number *K* (default: 10) functional expression programs *w*_*k*_. For each tree segment, we simulate the time evolution of expression programs by random walks with momentum term (see Fig. 1A and Supplementary Material). The mean expression of gene *g* in tree branch *b* at pseudotime *t* is a weighted sum of the *K* different programs 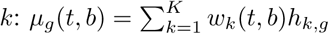 (Fig. 1B). The weights *h*_*k*, *g*_ are drawn from a gamma distribution (Supplementary Material, sections 6, 8).
3. **Sample cells from tree:** We offer three ways of sampling cells from a lineage tree, which impact the reconstruction difficulty: (1) sampling cells homogeneously along the tree, (2) sampling centered diffusely around selected tree points and (3) sampling with user-supplied density (Fig. 1C left, Supplementary Material, section 9).
4. **Simulate UMI counts:** We simulate unique molecular identifier (UMI) counts using a negative binomial distribution. First, a scaling factor *s*_*n*_ for the library size is drawn randomly for each cell *n* (see section 7 in Supplementary Material). Following Grün *et al.* (2014) and Harris *et al.* (2017), we make the variance 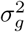 depend on the expected expression *s*_*n*_*μ*_*g*_ as 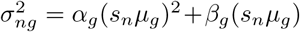. If x_*n*_(*t*, *b*) = (*x*_1_, *x*_2_, …, *x*_*G*_) is a cell at pseudotime *t* and branch *b*, the transcript counts are 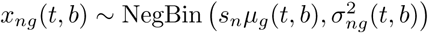 (Fig. 1C, right). For each of *N* cells and each of *G* genes we draw the number of UMIs from the negative binomial, resulting in an *N* × *G* expression matrix, which can serve as input for tree inference algorithms. Users can specify the topology of the lineage tree, assign branch pseudotime lengths, adjust parameters for the gene expression programs, and control the noise levels in the data. Default parameter values for *α*_*g*_, *β*_*g*_, and the base gene expression values were set in the range of parameters of real datasets (Supplementary Material, sections 3 and 8).

**Figure 1:**
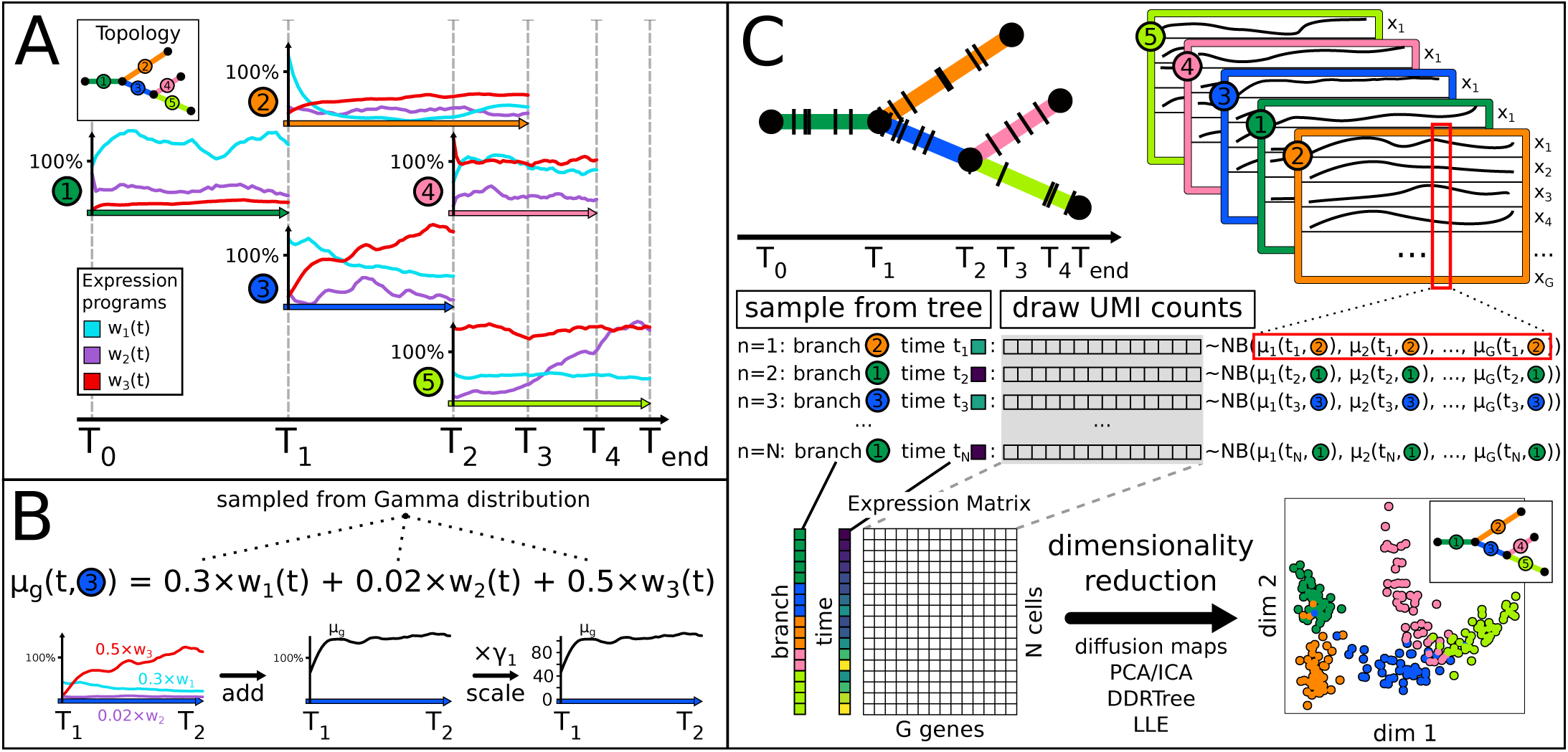
PROSSTT models the single-cell RNA-seq transcriptomes of cells differentiating along a (user supplied or sampled) lineage tree. (A) A small number of gene expression programs is simulated by random walk along each of the tree branches (number of steps = integer branch length). Here, a double bifurcation is regulated by thee expression programs. (B) Relative expected gene expression *μ*_*g*_(*t*, *b*) is computed as weighted sum of the expression programs with randomly sampled weights (here: gene *g* in branch 3). Expected expression values are obtained by multiplying with a gene-dependent sampled scaling factor. (C) Cells are sampled from the tree as pairs of pseudotime *t* and branch *b*. For each pair, the corresponding average gene expression is retrieved and UMI counts sampled using a negative binomial distribution. Low-dimensional representations of the resulting gene expression matrix are similar to those of real data (section 1, Supplementary Material) and capture the lineage tree topology (diffusion map created with destiny, (Angerer *et al.*, 2016)).

## 3 Application

We generated 10 sets of 100 simulations each, for different degrees of topology complexity (from 1 up to 10 bifurcations). In another study, we used this dataset to assess the performance of our tool MERLoT and other methods (Parra *et al.*, 2018).

## 4 Conclusions

PROSSTT simulates scRNA-seq data for complex differentiation processes. Low-dimensional visualizations produced by tree reconstruction tools resemble those of real datasets. Increasingly complex datasets with uncertain biological ground truth are becoming available. PROSSTT can help the development of methods that can reconstruct such complex trees by facilitating their quantitative assessment. Furthermore, the modular nature of the software allows for easy extensions, for example PROSSTT could serve to test the influence of noise models and give biological insights into how to model and interpret scRNA-seq data.

